# A dendritic-like microtubule network is organized from swellings of the basal fiber in neural progenitors

**DOI:** 10.1101/2020.03.16.993295

**Authors:** L. Coquand, G.S. Victoria, A. Tata, J.B. Brault, F. Guimiot, V. Fraisier, A. D. Baffet

## Abstract

Neurons of the neocortex are generated by neural progenitors called radial glial cells. These polarized cells extend a short apical process towards the ventricular surface and a long basal fiber that acts as a scaffold for neuronal migration. How the microtubule cytoskeleton is organized in these cells to support long-range transport in unknown. Using subcellular live imaging within brain tissue, we show that microtubules in the apical process uniformly emanate for the pericentrosomal region, while microtubules in the basal fiber display a mixed polarity, reminiscent of the mammalian dendrite. We identify acentrosomal microtubule organizing centers localized in swellings of the basal fiber. We characterize their distribution and demonstrate that they accumulate the minus end stabilizing factor CAMSAP3 and TGN-related membranes, from which the majority of microtubules grow. Finally, using live imaging of human fetal cortex, we show that this organization is conserved in basal radial glial (bRG) cells, a highly abundant progenitor cell population associated with human brain size expansion.

## Introduction

In the developing neocortex, neurons and glial cells are generated by neuronal progenitors called Radial Glial (RG) cells (Kriegstein and Alvarez-Buylla, 2009; Taverna et al., 2014). Two types of closely-related RG cells have been identified, with different localization, morphologies and abundancy. Apical radial glial (aRG) cells, also known as vRGs, are neuroepithelial cells present in all mammalian species. They are highly elongated bipolar cells, with an apical process attached to the ventricular surface, and a basal process (or fiber) extending towards the pial surface of the brain (Paridaen and Huttner, 2014) (Fig. 1A). Basal radial glial (bRG) cells, also known as oRGs, are rare in lissencephalic (smooth brain) species such as mice, and abundant in gyrencephalic (folded brain) species, including humans (Lui et al., 2011; Hansen et al., 2010; Fietz et al., 2010; Reillo et al., 2011). Their relative abundance is believed to account for variations in the size and degree of folding of the neocortex (Fernández et al., 2016). bRG cells derive from aRG but have delaminated from the neuroepithelium and retracted their apical process (Fig. 1A). aRG and bRG cells however share many characteristics, including a close transcriptional profile and an elongated basal process (Pollen et al., 2015; Hansen et al., 2010).

**Figure 1.**
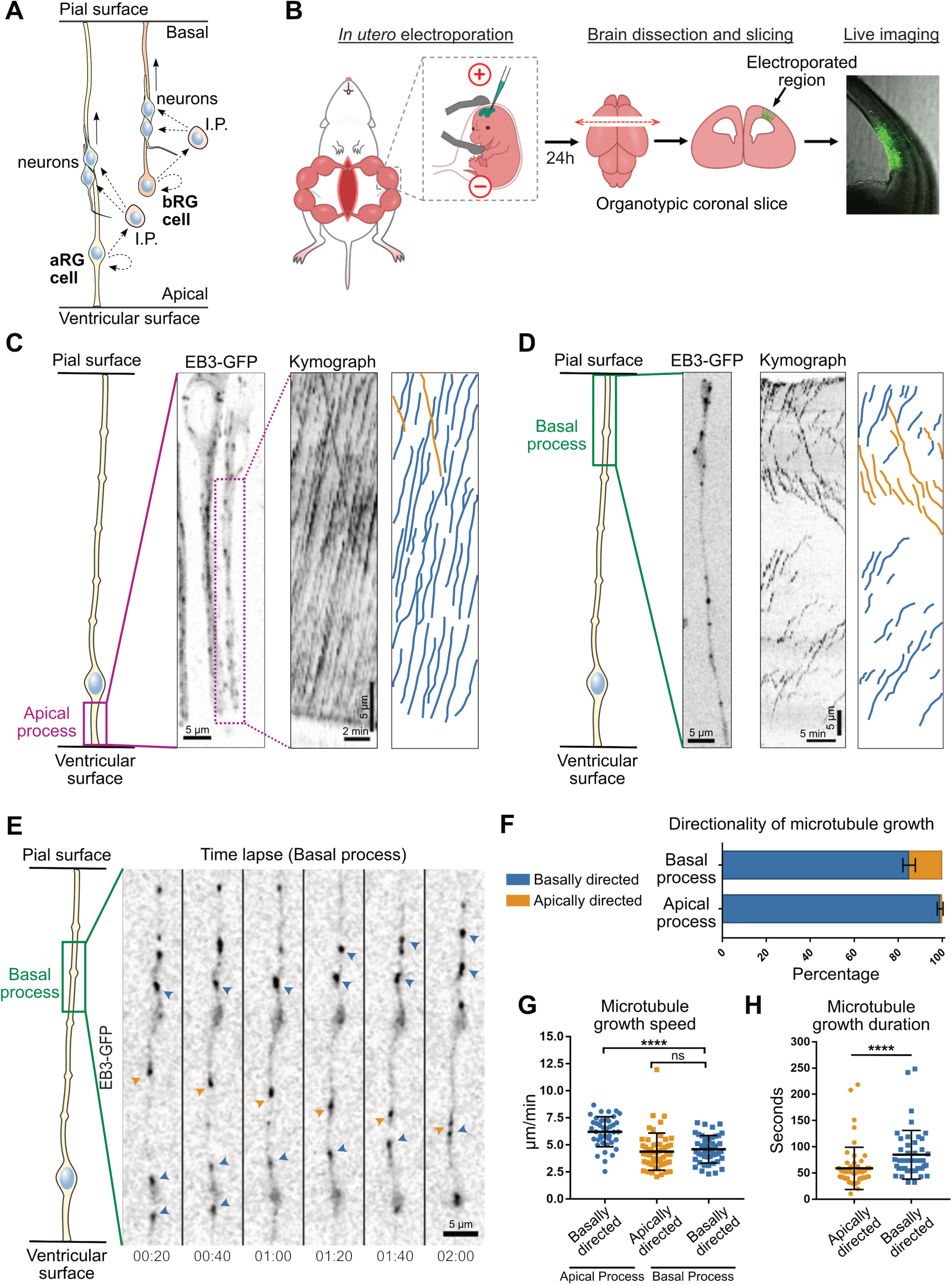
A bipolar microtubule network in the basal process of aRG cells. **A.** Schematic representation of apical radial glial (aRG) and basal radial glial (bRG) cells. These neural stem cells both generate neurons - via an intermediate progenitor (I.P.) - and support their migration, but bRG cells have lost their apical connection to the ventricular surface. As a convention, the basal side is represented upwards. **B.** Experimental set-up: Plasmids are injected into the left cortical ventricle of E13.5 embryos *in utero*. After 24h, the mother and embryos are sacrificed, embryonic cortices are recovered, sliced coronally and subjected to live imaging using high working distance objectives. **C.** (Left) Live imaging of EB3-GFP in the apical process of mouse aRG cells *in situ*. Scale bar = 5 *µ*m. (Center) Corresponding kymograph. Most comets originate from the apical endfoot and move basally. Scale bar = 2 min. (Right) Manual tracks corresponding to the kymograph. **D.** (Left) Live imaging of EB3-GFP in the basal process of mouse aRG cells *in situ*. Scale bar = 5 *µ*m. (Center) Corresponding kymograph. Comets display both apical and basal movement. Scale bar = 5 min. (Right) Manual tracks corresponding to the kymograph. **E.** Live imaging montage of EB3-GFP in the basal process of mouse aRG cell. Blue arrowheads indicate basally growing microtubules and orange arrowhead indicates apically growing microtubule. A swelling can be seen in the center. Scale bar = 5 *µ*m. **F.** Quantification of the average directionality of EB3 comets in the apical and basal processes (n = 302 and 827 comets, respectively). **G.** Quantification of the average speed of apically and basally directed EB3 comets in the basal process (n = 52 and 45 comets, respectively). **H.** Quantification of the average growth duration of apically and basally directed EB3 comets in the basal process (n = 52 and 45 comets, respectively). **F, G, H**, error bars indicate SD. ****p<0,0001 by Mann-Whitney tests.

The basal process of RG cells has long been known to act as a scaffold, guiding the migration of newborn neurons to their correct position in the neocortex (Noctor et al., 2004; Tan and Shi, 2013). More recently, the basal process has emerged as a potential regulator of cell fate (Shitamukai et al., 2011; Alexandre et al., 2010). Accordingly, a number of molecules important for basal process integrity or for RG cell proliferative capacity have been identified to localize in a polarized manner to the basal process (Yokota et al., 2009; 2010; Tsunekawa et al., 2012). A recent study identified multiple mRNAs localizing to the basal endfeet, and demonstrated their local translation (Pilaz et al., 2016). These mRNAs, bound to the RNA-binding protein Fmr1p, were shown to travel long distances within the basal process, at velocities consistent with microtubule-based transport.

Organization of the microtubule cytoskeleton is crucial for polarized transport of cargoes to various subcellular locations. While the centrosome is the main microtubule organizing center (MTOC) during mitosis, many differentiated cells - including neurons, myotubes or epithelial cells - display an acentrosomal microtubule organization during interphase (Bartolini and Gundersen, 2006). In neurons, the axonal microtubule network is unipolar with the plus ends pointing towards the axonal tip, while in dendrites, microtubules have a mixed polarity, with various amounts of “minus end out” microtubules, depending on the neuronal type and species (Yau et al., 2016; Baas et al., 1988). This particular microtubule organization depends on γ-tubulin-mediated acentrosomal nucleation, as well as on CAMSAP/Patronin-mediated minus end growth (Ori-McKenney et al., 2012; Wang et al., 2019; Feng et al., 2019; Pongrakhananon et al., 2018; Yau et al., 2014; Marcette et al., 2014; Chuang et al., 2014).

A variety of genetic mutations have been shown to lead to malformations of cortical development (MCDs), such as microcephaly and lissencephaly (Pinson et al., 2019). Strikingly, the majority of affected genes code for proteins associated with the microtubule cytoskeleton (Poirier et al., 2013; Jayaraman et al., 2018; Reiner et al., 1993). Few regulators of microtubule organization have been investigated so far in RG cells. The adaptor protein Memo1 controls the localization of CAMSAP2 and the stability and organization of the microtubule network in dissociated mouse RG cell cultures (Nakagawa et al., 2019). In the apical process, the centrosomal protein AKNA promotes microtubule nucleation and regulates aRG cell delamination (Camargo Ortega et al., 2019). The organization and polarity of the microtubule network in aRG and bRG cells *in situ* is however currently unknown. This is largely due to the challenge of studying dynamic subcellular processes in real time within thick organotypic brain cultures.

Here, using an approach for subcellular live imagining within cerebral tissue, we characterize microtubule organization in mouse aRG and human bRG cells *in situ*, within the native architecture of the cortex. We determine that, while microtubule polarity in the apical process of mouse aRG cells is unipolar, the basal process displays a mixed microtubule polarity, reminiscent of dendritic arbors in vertebrate neurons. We further identify acentrosomal microtubule organizing centers localized in swellings of the basal process and, using live imaging of human fetal brain slices, demonstrate that this organization is a conserved feature of human bRG cells.

## Results

### A bipolar microtubule network in the basal process of aRG cells

To visualize the orientation of growing microtubules in mouse aRG cells *in situ*, we developed an approach for high resolution and fast subcellular live imaging within thick embryonic brain slices. GFP-tagged plus-end tracking protein (+TIP) EB3 was delivered to aRG cells using *in utero* electroporation at embryonic day 13.5 (E13.5) (Baffet et al., 2016). The embryos were then sacrificed 24h later, and brains were sliced and mounted for imaging using a modified sample preparation and imaging method (see methods) (Fig. 1B). We first revisited microtubule organization in the apical process of mouse aRG cells. This analysis indicated that over 99% of microtubule plus ends emanated from the apical endfoot, where the centrosome is located, and grew in the basal direction towards the cell soma (Fig. 1C, 1F & Supplemental Movie 1). We then performed a similar analysis in the basal process of aRG cells. In contrast to what we observed in the apical process, growing microtubules adopted a mixed polarity, reminiscent of dendritic microtubule organization (Fig. 1D, 1E & Supplemental Movie 2 and 3). This organization, however, remained biased towards basally-directed growth, as only 15% of microtubules grew in the apical direction (Fig. 1F). Apically and basally-directed microtubules within the basal fiber grew at similar speeds, but slower than in the apical process (Fig. 1G). In the basal process, basally-directed microtubules grew for longer durations, suggesting higher stability (Fig. 1H). EB3 comets did not grow from the apical centrosome, which is located hundreds of *µ*m away, but directly emanated from the basal process. Therefore, microtubules in the apical process of aRG cells emanate from the pericentrosomal region and form a unipolar network growing in the basal direction, while microtubules in the basal process appear largely acentrosomal, and have a bipolar orientation biased towards basal growth.

### Acentrosomal microtubules preferentially grow from swellings of the basal process

We next asked whether acentrosomal microtubule organizing centers (aMTOCs) may exist within the basal process of aRG cells. From the observation of the EB3-GFP movies, we noted that a large number of newly appearing comets emanated from swellings of the basal process (Fig 1D). Swellings (also known as varicosities) are well-known but poorly-described deformations of RG cell basal processes, with no reported function (Noctor et al., 2001; Hansen et al., 2010; Hu et al., 2013). We could easily observe these structures following expression of soluble GFP or immuno-staining against the radial glial-specific protein Nestin, allowing visualization of the RG cell outline (Fig. 2A, 2B). To measure microtubule growth from these structures, we live imaged a large number of swellings as well as basal process shafts (non-deformed regions) in EB3-GFP-expressing cells, and quantified the rate of new EB3 comet formation in these two domains. This analysis revealed that the average rate of comet formation in swellings was 7,09 times higher than in the rest of the shaft (Fig. 2C, 2D & Supplemental Movie 4). Moreover, while 89,6% of comets emanating from the shaft grew in the basal direction, microtubules emanating from swellings appeared initially more randomly oriented, albeit still with a basal preference (65,7%) (Fig. 2E). This analysis therefore identifies swellings of the basal process of mouse aRG cells as acentrosomal microtubule organizing centers.

**Figure 2.**
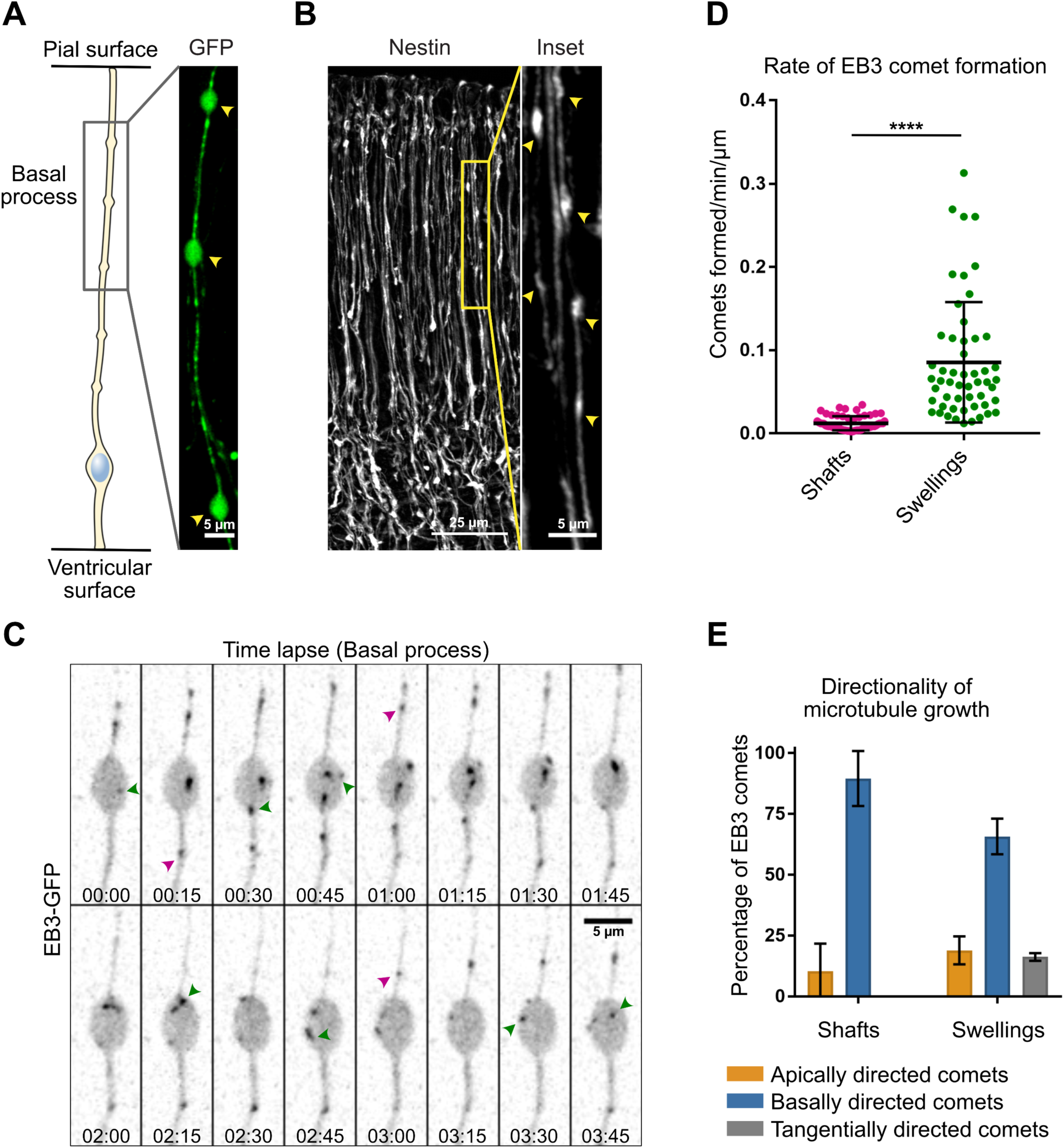
Acentrosomal microtubules preferentially grow from swellings of the basal process. **A, B.** The basal processes of aRG cells display swellings along their length, which can be visualized by overexpression of GFP (**A**, scale bar = 5 *µ*m) as well as by immunofluorescence against Nestin, a RG cell-specific cytoskeletal marker (**B**, scale bar = 25 *µ*m. Inset, scale bar = 5 *µ*m). **C.** Live imaging of EB3-GFP in the basal process of an aRG cell showing the emergence of new comets within a swelling. Green arrowheads: newborn comets in the swelling. Purple arrowheads: newborn comets in the shaft. Scale bar = 5 *µ*m. **D.** Quantification of the rate of EB3 comet formation in basal process shafts and swellings, normalized to length (n=52 shafts and 53 swellings). **E.** Quantification of the average directionality of EB3 comets in the shafts and swellings of the basal processes (n = 122 and 260 comets, respectively). **D, E**, error bars indicate SD. ****p<0,0001 by Mann-Whitney tests.

### Dendritic-like microtubules organization from swellings is a conserved feature of human bRG cells

Because bRG cells share many characteristics with aRG cells, we next asked if bipolar microtubule organization from basal process swellings was also a feature of human bRG cells. We reasoned that this may be even more critical for these cells, which can be millimeters long in the human brain. To test this, we developed a protocol to electroporate and live image pieces of human fetal frontal cortex biopsies, obtained from second trimester medical pregnancy terminations (Fig. 3A, 3B & see Methods). We identified bRG cells based on their localization in the subventricular zone and morphology. Their identity was confirmed after performing immuno-staining against SOX2, a RG cell marker. We first confirmed the presence of numerous swellings all along the basal process of human bRG cells, which were visible following GFP electroporation or immuno-staining against Vimentin (Fig. 3C, 3D). We next expressed EB3-GFP in human fetal cortex samples and recorded plus end microtubule growth in bRG cell swellings. Similar to our observations in mouse aRG cells, we observed abundant *de novo* EB3 comet formation within swellings (Fig. 3E & Supplemental Movie 5). The rate of EB3 comet formation inside swellings was very similar in the two cell types (Fig. 3F). Finally, we analyzed the directionality of EB3 comets in the basal process of human bRG cells, which revealed, as for mouse aRG cells, a bipolar microtubule network biased towards basal growth (82,3 ± 84,5 %) (Fig. 3G). Therefore, bipolar microtubule network organization from basal process swellings appears to be conserved between mouse aRG and human bRG neural stem cells.

**Figure 3.**
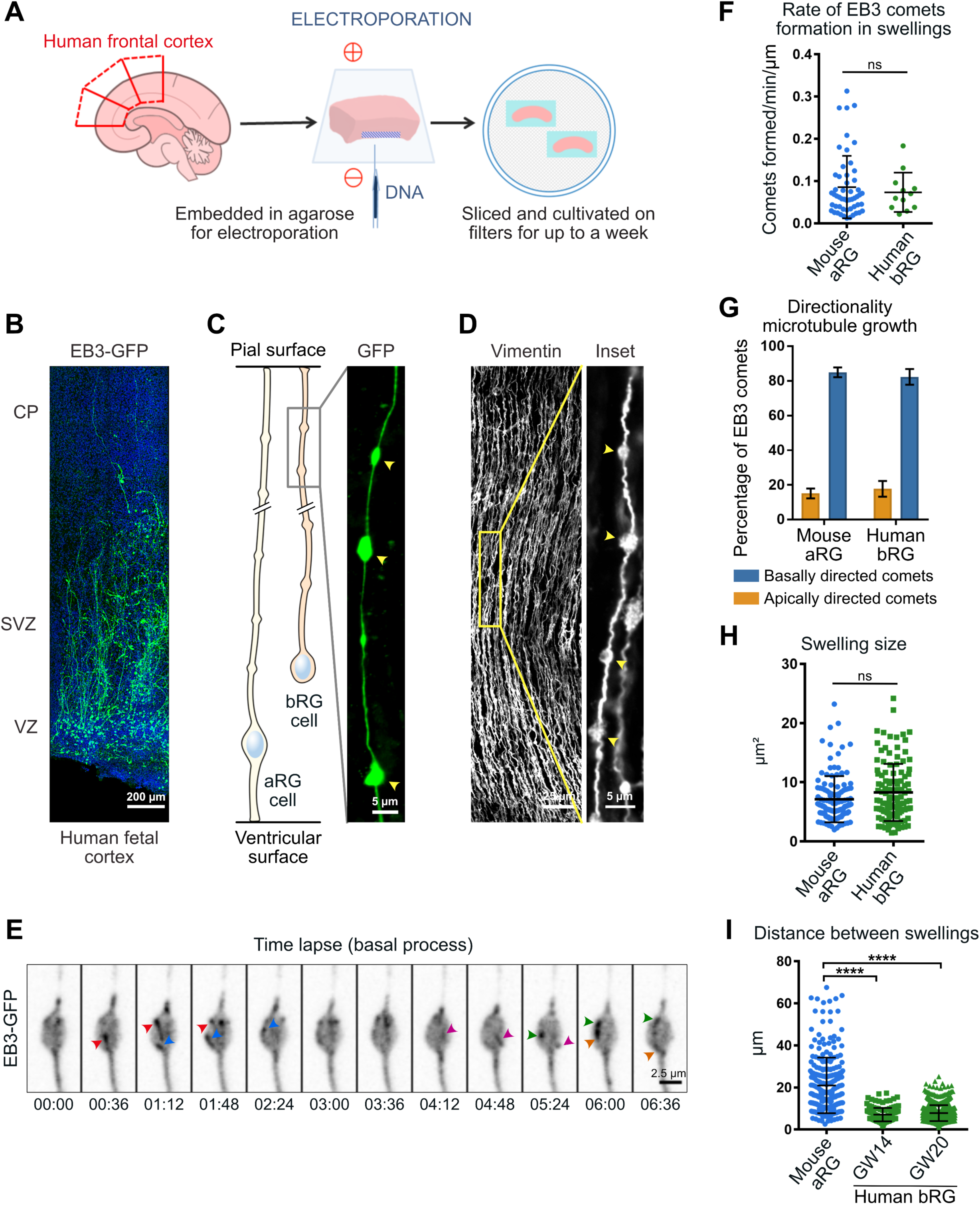
Dendritic-like microtubules organization from swellings is a conserved feature of human bRG cells. **A.** Schematic of protocol as described in Methods. Pieces of human foetal cortex were embedded in agarose prior to injection with DNA underneath the ventricular zone (VZ) and electroporation, followed by slicing and culture. **B.** A slice of human foetal cortex cultured for 72 hours *in vitro* prior to fixation and staining with DAPI (blue). EB3-GFP-electroporated basal radial glia (green) occupy the subventricular zone (SVZ) and extend long basal processes. VZ: Ventricular Zone, CP: Cortical plate. Scale bar: 200 *µ*m. **C, D.** The basal processes of human bRG cells display swellings along their length, which can be visualized by overexpression of GFP (**C**, scale bar = 5 *µ*m) as well as by immunofluorescence against Vimentin, a RG cell-specific cytoskeletal marker (**D**, scale bar = 25 *µ*m. Inset, scale bar = 5 *µ*m). **E.** Live imaging of a human bRG cell showing appearance of *de novo* EB3 comets (arrowheads) in a basal process swelling. Scale bar: 5 *µ*m. **F.** Quantification of the rate of EB3 comet formation in basal process swellings of human bRG cells at gestational week (GW) 18. Mouse data from figure 2D are shown for comparison. (n=12 human swellings and 53 mouse swellings) **G.** Quantification of the average directionality of EB3 comets in the basal process of human bRG cells (n = 205 comets). Mouse data from figure 1F are shown for comparison. **H.** Quantification of swelling size in mouse (E14.5) and human (GW 14 and 20) tissue (n=55, 129 and 772 swellings respectively). **I.** Quantification of the distance between individual swellings along the basal process in mouse (E14.5) and human (GW 14 and 20) tissue (n=260 and 605 swellings respectively). In human tissue, swellings occur with greater frequency but are constant between early and late neurogenic stages. **F, G, H, I**, error bars indicate SD. ****p<0,0001 by Mann-Whitney tests.

### Size and distribution of basal process swellings

While the role of basal process swellings remains unexplored, a description of their morphology and periodicity is also lacking. We first compared the size of swellings in mouse aRG and human fetal bRG cells. This analysis revealed relatively variable sizes, but on average similar between the two cell types (around 8 *µ*m^2^ for mouse aRG cells and 9 *µ*m^2^ for human bRG cells) (Fig. 3H). We next analyzed the distribution of swellings along the basal process by measuring the average swelling-to-swelling distance. While the distance between two consecutive swellings was quite variable, their frequency was substantially higher in human bRG cells than in mouse aRG cells (1 every 7,7 *µ*m vs 1 every 21 *µ*m, respectively) (Fig. 3I). The higher frequency of swellings in bRG cells may be due to their greater length, requiring more microtubule organizing centers far away from the centrosome. However, human swelling distribution in early (gestational week 14) and late neurogenic stage - when the basal process is substantially longer (week 20) - remained constant (Fig 3I).

### Basal process microtubules grow from CAMSAP3-positive foci

To identify how swellings may act as local acentrosomal MTOCs, we first investigated the localization of the key microtubule nucleator, the γ-tubulin ring complex (γ-TURC). Expression of mEmerald-γ-Tubulin revealed its expected enrichment in the pericentrosomal region at the base of the apical process (Fig. 4A). However, γ-tubulin was undetectable within the basal process, both in swellings or in the rest of the shaft (Fig. 4A), suggesting an absence of γ-TURC-based nucleation away from the centrosome. We next tested whether growing microtubules preferentially emerged from basal process swellings due to an accumulation of stabilized microtubule minus ends within these structures. CAMSAP3 specifically recognizes and stabilizes microtubule minus ends, generating seeds from which multiple rounds of plus end growth and shrinkage can occur (Jiang et al., 2014). Upon electroporation, GFP-CAMSAP3 accumulated not only at the ventricular surface but, unlike γ-tubulin, also localized as patches along the basal process of aRG cells (Fig. 4B, 4C). CAMSAP3 could be observed inside as well as outside swellings, consistent with EB3 comets appearing in both locations (Fig. 4C). However, the higher frequency of EB3 growth from swellings was reflected by the very high percentage of swellings containing CAMSAP3 clusters (85,1 ± 8,4 %) (Fig. 4D). Moreover, CAMSAP3 clusters were much larger within swellings than in the shaft (Fig. 4C).

**Figure 4.**
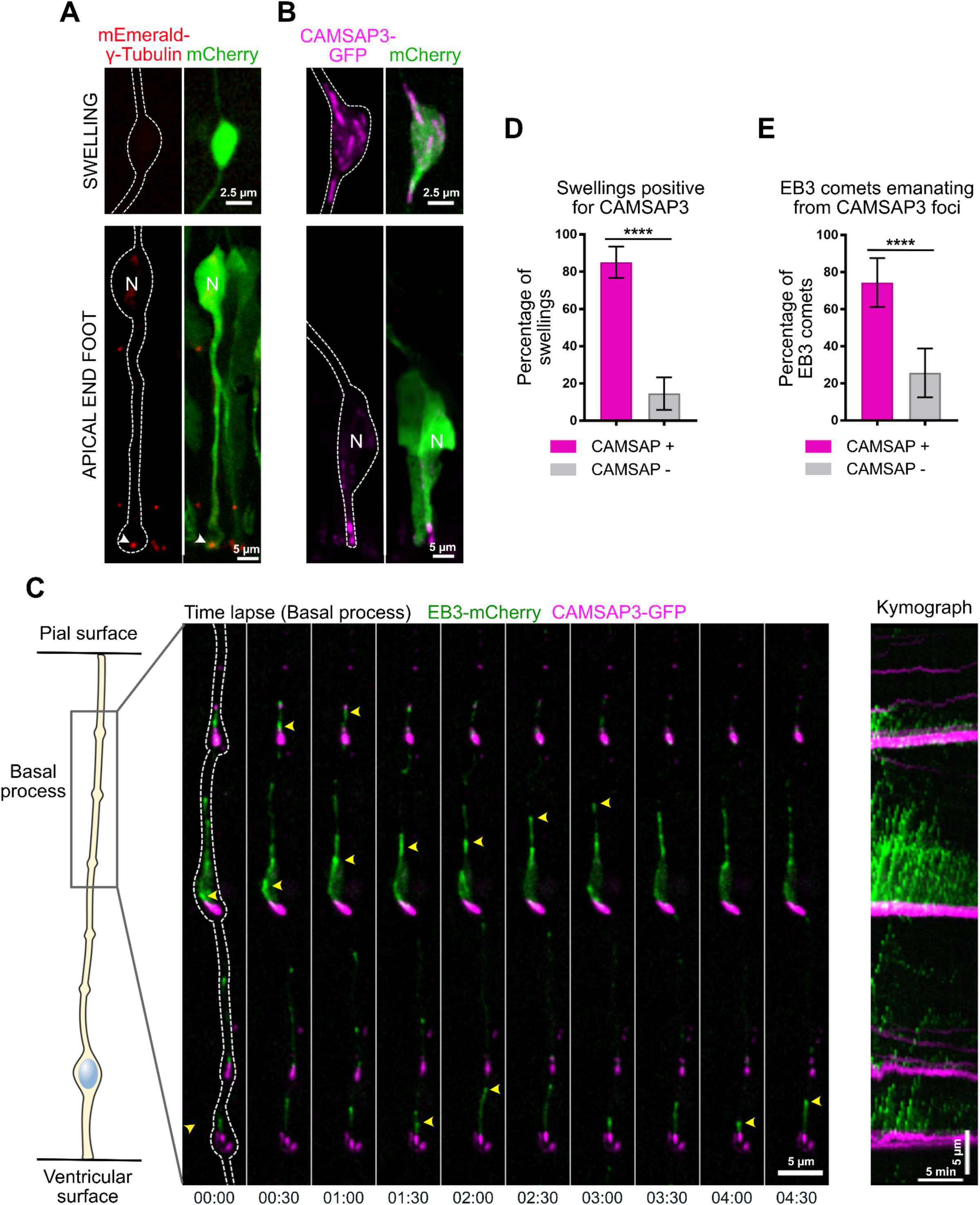
Basal process microtubules grow from CAMSAP3-positive foci. **A, B.** mEmerald-γ-tubulin and CAMSAP3-GFP localization in mouse aRG cell apical endfeet (scale bar = 5 *µ*m) and basal process swellings (scale bar = 2,5 *µ*m). Co-expression of mCherry allows to visualize RG cell outline. Both γ-tubulin and CAMSAP3 localize to the apical surface, but only CAMSAP3 localizes to basal process swellings. **C.** (Left) Live imaging of CAMSAP3-GFP and EB3-mCherry in mouse aRG cell basal process. EB3 comets emanate from CAMSAP3-positive foci, which are concentrated within basal process swellings. Scale bar = 5 *µ*m. (Right) Corresponding kymograph. Scale bar = 5 minutes. **D.** Quantification of the percentage of basal process swellings positive for CAMSAP3 (n=120 swellings). **E.** Quantification of the percentage of EB3 comets emanating from CAMSAP3-positive foci (n = 345 comets). ****p<0,0001 by Mann-Whitney tests.

We then live imaged GFP-CAMSAP3 together with EB3-mCherry, within embryonic brain slices. In contrast to the highly dynamic EB3 comets, CAMSAP3 foci remained relatively immobile, as expected for stabilized microtubule minus ends. The majority of newly formed EB3-mCherry comets were observed emanating from CAMSAP3-GFP clusters (74.4 ± 13.2%) (Fig. 4C, 4E & Supplemental Movie 6). This was the case within swellings, but also in the shaft where the less frequent formation of novel EB3 comets still strongly correlated with CAMSAP3 foci. These results suggest that microtubules preferentially grow from stabilized minus ends concentrated within swellings of the basal process.

### A TGN-related compartment localizes to microtubule minus ends in basal process swellings

Because the Golgi apparatus is a major site for microtubule organization, we next asked whether Golgi outposts could be found along the basal process of aRG cells, similarly to what happens in neurons (Ori-McKenney et al., 2012). To test this, we *in utero* electroporated the cerebral cortex of mouse embryos with constructs expressing different tagged Golgi markers along with a cytoplasmic fluorescent protein to visualize basal process outline. In the apical process, where the Golgi apparatus is localized, we consistently detected the *cis-medial* marker ManII, the *trans* marker GalNAcT2, as well as the small GTPase Rab6A (Fig. 5A). Strikingly, the *trans* and Trans Golgi Network (TGN) markers Rab6A, GalNAcT2, GalT and TGN46 also localized within the basal process and accumulated within the vast majority of swellings (87,8 ± 5,1% for GalNAcT2) (Fig. 5B, 5E). The *cis-medial* markers ManII and GMAP210 were, however, undetectable outside the apical process (Fig. 5B). These results point towards the presence of a Golgi-related secretory compartment with a *trans-*Golgi*/*TGN identity in basal process swellings of RG cells.

**Figure 5.**
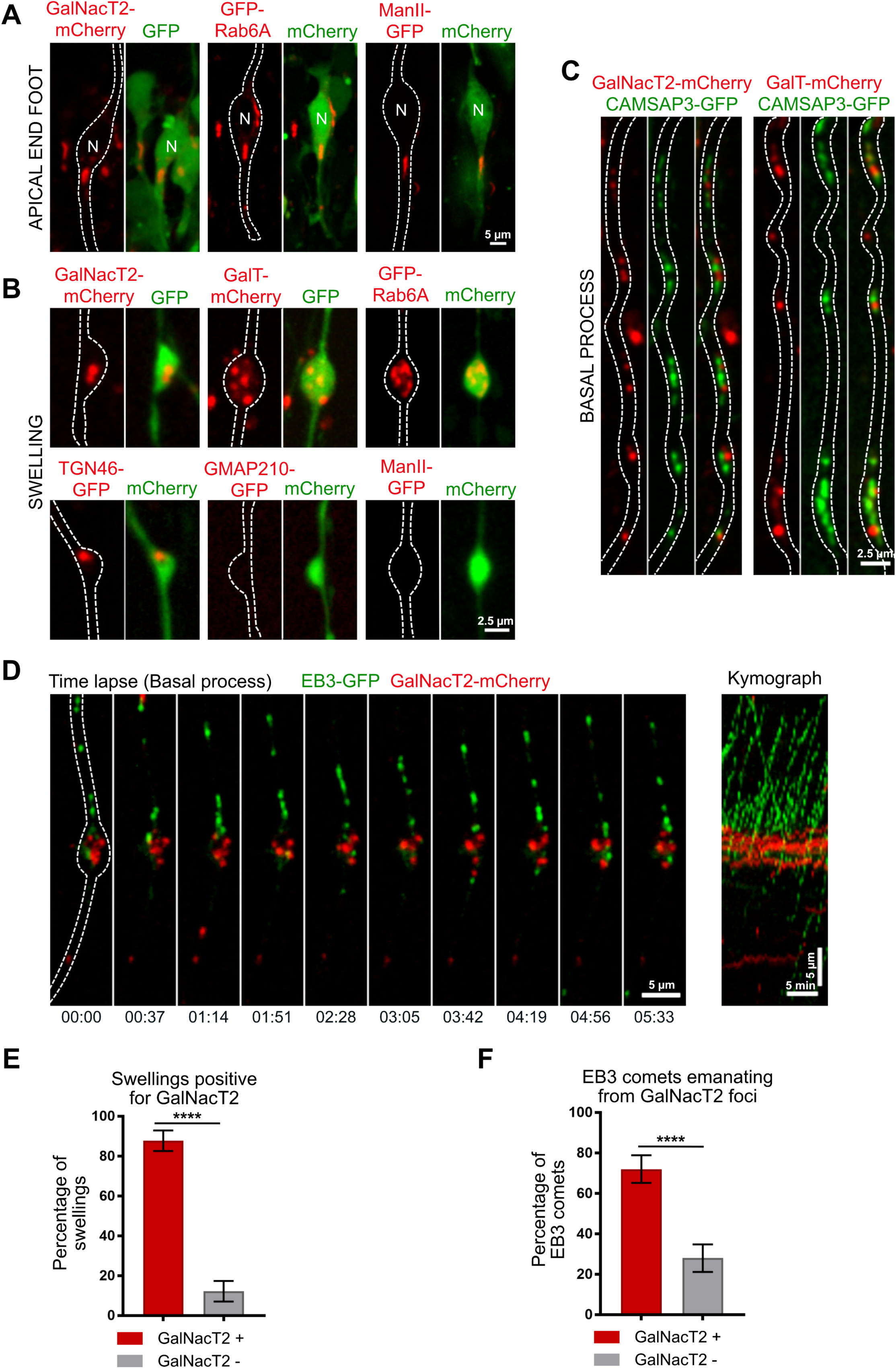
A Golgi-related compartment localizes to microtubule minus ends in basal process swellings. **A.** Expression of GalNacT2-mCherry, GFP-Rab6A and ManII-GFP in aRG cells highlighting localization of the Golgi apparatus in the apical process, next to the nucleus but away from the apical surface. scale bar = 5 *µ*m **B.** Expression of GalNacT2-mCherry, GalT-mCherry, GFP-RAB6, TGN46-GFP, GMAP210-GFP and ManII-GFP in aRG cells revealing localization of GalNacT2, GalT, RAB6, TGN46, and absence of GMAP210 and ManII, in basal process swellings. scale bar = 2,5 *µ*m. **A, B**, GFP or mCherry co-expression allows visualization of cell outline. **C.** Co-expression of GalNacT2-mCherry and GalT-mCherry with CAMSAP3-GFP reveals a close apposition along the basal process of mouse aRG cells. Scale bar = 2,5 *µ*m. **D.** (Left) Live imaging of EB3-GFP and GalNacT2-mCherry in mouse aRG cell basal process. EB3 comets emanate from GalNacT2-positive foci. Scale bar = 5 *µ*m. (Right) Corresponding kymograph. Scale bar = 5 minutes. **F.** Quantification of the percentage of basal process swellings positive for GalNacT2 (n = 70 swellings). **F.** Quantification of the percentage of EB3 comets emanating from GalNacT2-positive foci (n = 333 comets). ****p<0,0001 by Mann-Whitney tests.

CAMSAP2 was previously shown to be specifically recruited to the *cis*-Golgi (Wu et al., 2016). Consistently, we noted that its homologue CAMSAP3 did not colocalize with the *trans* marker GalNacT2 in basal process swellings (Fig. 5C). However, we noted that CAMSAP3 and GalNacT2 were frequently found in close proximity (Fig. 5C). Because the *trans* and TGN can stimulate microtubule growth, via the recruitment of CLASP 1 and 2 (Efimov et al., 2007), we asked whether microtubules preferentially grew from GalNacT2-positive foci. We therefore live imaged GalNAcT2 together with EB3, and quantified the amount of newly formed EB3 comets emanating for GalNAcT2-positive clusters. In agreement with the CAMSAP3-GalNacT2 apposition, this analysis revealed a strong association between GalNacT2-positive structures and newly formed EB3 comets (72 ± 6.8%) (Fig. 5D, 5F & Supplemental Movie 7). These results point towards a potential role for Golgi membranes in the growth of minus end-stabilized microtubules, within swellings of RG cell basal processes.

## DISCUSSION

In this report, we characterize the microtubule network organization in radial glial cells, the neural stem cells of the developing mammalian neocortex. We show that microtubules of the basal process are acentrosomal and originate predominantly from swellings where we find Golgi-related membranes as well as the minus end stabilizing factor CAMSAP3 to accumulate. Moreover, we demonstrate that this organization is conserved from mouse aRG cells to human bRG cells. Unlike for neurons, which still polarize when cultivated *in vitro*, analysis of RG cells can only be performed within the tissue. Indeed, RG cells cultivated *in vitro* lose their apico-basal polarity and do not generate any swelling. Our study was therefore entirely performed *in situ*, within thick embryonic brain slices.

The basal process organization is reminiscent of what has been described in the mammalian dendrite, where one third of dynamic microtubules are “minus end out”, growing towards the soma (Baas et al., 1988; Yau et al., 2016). This polarity suggests that trafficking into the basal process is likely to rely on kinesin-based movement, but that minus end-based transport of specific cargos – as shown in dendrites (Kapitein et al., 2010) – cannot be ruled out. It is important to stress that EB3-tracking allows to measure the polarity of the dynamic microtubule network, and is only an approximation for the overall microtubule polarity. Indeed, laser-cut and motor-PAINT experiments performed in hippocampal neurons in culture have revealed an even greater proportion of “minus end out” microtubules (Yau et al., 2016; Tas et al., 2017). The identification of microtubule organizing centers throughout the basal process is consistent with microtubules growing both apically and basally, but how the polarity of the network is biased towards basal growth remains unclear. In axons, unipolar microtubule organization depends on the HAUS/augmin complex (Cunha-Ferreira et al., 2018; Sánchez-Huertas et al., 2016). This complex was recently shown to be critical for γ-tubulin-mediated nucleation from presynaptic boutons (Qu et al., 2019), where increased microtubule dynamics favors delivery of synaptic vesicle precursors (Guedes-Dias et al., 2019).

Based on the absence of γ-tubulin, swellings do not appear to be sites of microtubule nucleation, but rather sites of minus end stabilization. Although we cannot rule out the presence of undetectable amounts of γ-tubulin in swellings, the localization of the minus end capping-protein CAMSAP3 further supports this notion. If microtubules are not nucleated within the basal process, where could stabilized seeds come from? One possibility is that acentrosomal microtubules are generated in the apical process by severing enzymes such as spastin or katanin. Alternatively, such severing could occur directly within the basal process, in order to amplify the number of acentrosomal microtubules. Interestingly, human mutations in *KATNB1*, which encodes the p80 subunit of katanin, cause severe microcephaly and lissencephaly (Hu et al., 2014). While mitotic and ciliary defects were reported in mutant RG cells, the role of katanin in interphasic microtubule organization was not addressed.

We observed the presence of *trans* and TGN markers, but not *cis-medial* elements, in swellings of the basal process. This argues against the presence of canonical Golgi outposts, and rather points to a secretory structure, presumably with a TGN identity. The TGN is known to locally stimulate microtubule growth via recruitment of the CLASPs plus end binding proteins (Efimov et al., 2007). The presence of glycosylating enzymes (i.e. GalT) is surprising and may reflect leakage from the apical Golgi apparatus. Nevertheless, our observations are consistent with an electron microscopy study revealing a lack of Golgi cisternae in the basal process of aRG cells (Taverna et al., 2016). Our data therefore argue against Golgi-mediated nucleation, but suggest a potential role for TGN membranes in microtubule plus end growth. Together, this work identifies the organization of the microtubule cytoskeleton in mouse and human RG cells, and will allow to determine how genetic mutations targeting microtubule regulators may affect these neural progenitor cells, leading to brain malformations.

## Materials and Methods

### Ethics statement

For animal care, we followed the European and French National Regulation for the Protection of Vertebrate Animals used for Experimental and other Scientific Purposes (Directive 2010/63; French Decree 2013-118). The project was authorized and benefited from guidance of the Animal Welfare Body, Research Centre, Institut Curie. CD1-IGS pregnant females were purchased from Charles River Laboratories (France).

Human fetal tissue samples were collected with previous patient consent and in strict observance of legal and institutional ethical regulations. The protocol was approved by the French biomedical agency (Agence de la Biomédecine, approval number: PFS17-003).

### *In utero* electroporation of mouse embryonic cortex

Pregnant CD1-IGS mice at embryonic day 13.5 (E13.5) were anesthetized with isoflurane gas, and injected subcutaneously first with buprenorphine (0.075mg/kg) and a local analgesic, bupivacaine (2 mg/kg), at the site of the incision. Lacrinorm gel was applied to the eyes to prevent dryness/irritation during the surgery. The abdomen was shaved and disinfected with ethanol and antibiotic swabs, then opened, and the uterine horns exposed. Plasmid DNA mixtures were used at a final concentration of 1 *µ*g/*µ*l per plasmid, dyed with Fast Green and injected into the left lateral ventricle of several embryos. The embryos were then electroporated through the uterine walls with a NEPA21 Electroporator (Nepagene) and a platinum plated electrode (5 pulses of 50 V for 50 ms at 1 second intervals). The uterus was replaced and the abdomen sutured. The mother was allowed to recover from surgery and supplied with painkillers in drinking water post-surgery.

### Electroporation of human fetal cortex

Fresh tissue from human fetal cortex was obtained from autopsies. A piece of pre-frontal cortex was collected from one hemisphere, and transported on ice to the lab. The tissue was divided into smaller pieces and embedded 4% low-gelling agarose (Sigma) dissolved in artificial cerebrospinal fluid (ACSF). Plasmid DNA (1 *µ*g/*µ*l) was injected with a fine glass micropipette through the agarose at the ventricular surface. The gel block was then subjected to a series of 5 pulses of 50 V for 50 ms at 1 second intervals and sliced with a Leica VT1200S vibratome (300 *µ*m-thick slices) in ice-cold ACSF. Slices were grown on Millicell culture inserts (Merck) in cortical culture medium (DMEM-F12 containing B27, N2, 10 ng/ml FGF, 10 ng/ml EGF, 5% fetal bovine serum and 5% horse serum) for up to 5 days. Medium was changed every day.

### Subcellular live imaging in mouse embryonic brain and human fetal cortex slices

To record EB3-GFP or EB3-mCherry dynamics together with Golgi markers or CAMSAP3-GFP in radial glia *in situ*, we used the following approach. 24h after the electroporation, the pregnant mouse was sacrificed and the electroporated embryos recovered. Brains were dissected in ACSF and 300 *µ*m-thick coronal slices were prepared with a Leica VT1200S vibratome in ice-cold ACSF. The slices were cultured on membrane filters over enriched medium (DMEM-F12 containing B27, N2, 10 ng/ml FGF, 10 ng/ml EGF, 5% fetal bovine serum and 5% horse serum). After recovery in an incubator at 37°C, 5% CO2 for 2 hours (or 24H for human tissue), the filters were cut and carefully turned over on a 35 mm FluoroDish (WPI), in order to position the sample in direct contact with the glass, underneath the filter (which maintained the sample flat). Live imaging was performed on a spinning disk wide microscope equipped with a Yokogawa CSU-W1 scanner unit to increase the field of view and improve the resolution deep in the sample. The microscope was equipped with a high working distance (WD 0.3 mm) 100X SR HP Plan Apo 1.35 NA Silicon immersion (Nikon), or a 60X 1.27 NA Apo plan objective (Nikon), and a Prime95B SCMOS camera. Z-stacks of 20-30 *µ*m range were taken with an interval of 1 *µ*m. Videos were mounted in Metamorph. Image analysis (Kymographs and other quantifications), modifications of brightness and contrast were carried out on Fiji. Figures were assembled in Affinity Designer.

### Immunostaining of brain slices

Mouse embryonic brains were dissected out of the skull, fixed in 4% Pfa for 2 hours, and 80 *µ*m-thick slices were prepared with a Leica VT1200S vibratome in PBS. Human fetal slices in culture were fixed in 4% Pfa for 2 hours. Slices were boiled in citrate sodium buffer (10mM, pH6) for 20 minutes and cooled down at room temperature (antigen retrieval). Slices were then blocked in PBS-Triton X100 0.3%-Donkey serum 2% at room temperature for 2 hours, incubated with primary antibody overnight at 4°C in blocking solution, washed in PBS-Tween 0.05%, and incubated with secondary antibody overnight at 4°C in blocking solution before final wash and mounting in aquapolymount. Mosaics (tilescans) of fixed human tissue were acquired with a 40X Apo-Plan objective.

### Expression constructs and antibodies

The following plasmids were used in this study. CAMSAP3-GFP (a gift from Masatoshi Takeichi); EB3-GFP (a gift from Matthieu Piel); EB3-mCherry (Michael Davidson, Addgene 55037); mCherry2-C1 empty vector (Michael Davidson, Addgene 54563); mEGFP-C1 empty vector (Michael Davidson, Addgene 54759); mEmerald-γ-Tubulin (Michael Davidson, Addgene 54105); GFP-Rab6A (a gift from Bruno Goud); GFP-GMAP210 (a gift from Claire Hivroz); GalT-mCherry, GalNacT2-mCherry, ManII-GFP, TGN46-GFP (all gifts from Franck Perez). Antibodies used in this study were mouse anti-Nestin (BD Pharmingen 556309, 1/500) and rat anti-Vimentin (R&D systems MAB2105, 1/500).

## Acknowledgments

We acknowledge the Cell and Tissue Imaging (PICT-IBiSA), Institut Curie, member of the French National Research Infrastructure France-BioImaging (ANR10-INBS-04) and the Nikon BioImaging Center (Institut Curie, France). We thank F. Perez, Matthieu Piel, Claire Hivroz, Bruno Goud (I. Curie) and Masatoshi Takeichi (Riken) for reagents and advice. We thank Renata Basto, Franck Perez and Bruno Goud (I. Curie) for helpful discussions and critical reading of the manuscript. L.C. was funded by a PSL/Sorbonne Université fellowship, G.S.V. by I. Curie and the Ville de Paris, J.B.B. by PSL and the Ville de Paris. A.D.B. is an Inserm researcher. This work was supported by the CNRS, I. Curie, the Ville de Paris “Emergences” program, Labex CelTisPhyBio (11-LBX-0038) and PSL.

## Author contributions

L.C., G.S.V. and A.D.B. conceived the project, analysed the data and wrote the manuscript. L.C. and G.S.V. did most of the experimental procedures. A.T. performed swelling characterization experiments. J.B.B and V.F. contributed with high resolution microscopy in the brain. F.G. provided human foetal sample. A.D.B. supervised the project.

## References

Alexandre, P., A.M. Reugels, D. Barker, E. Blanc, and J.D.W. Clarke. 2010. Neurons derive from the more apical daughter in asymmetric divisions in the zebrafish neural tube. Nature Publishing Group. 13:673–679. doi:10.1038/nn.2547.

Baas, P.W., J.S. Deitch, M.M. Black, and G.A. Banker. 1988. Polarity orientation of microtubules in hippocampal neurons: uniformity in the axon and nonuniformity in the dendrite. Proc Natl Acad Sci USA. 85:8335–8339. doi:10.1073/pnas.85.21.8335.

Baffet, A.D., A. Carabalona, T.J. Dantas, D.D. Doobin, D.J. Hu, and R.B. Vallee. 2016. Cellular and subcellular imaging of motor protein-based behavior in embryonic rat brain. Methods Cell Biol. 131:349–363. doi:10.1016/bs.mcb.2015.06.013.

Bartolini, F., and G.G. Gundersen. 2006. Generation of noncentrosomal microtubule arrays. J Cell Sci. 119:4155–4163. doi:10.1242/jcs.03227.

Camargo Ortega, G., S. Falk, P.A. Johansson, E. Peyre, L. Broix, S.K. Sahu, W. Hirst, T. Schlichthaerle, C. De Juan Romero, K. Draganova, S. Vinopal, K. Chinnappa, A. Gavranovic, T. Karakaya, T. Steininger, J. Merl-Pham, R. Feederle, W. Shao, S.-H. Shi, S.M. Hauck, R. Jungmann, F. Bradke, V. Borrell, A. Geerlof, S. Reber, V.K. Tiwari, W.B. Huttner, M. Wilsch-Bräuninger, L. Nguyen, and M. Götz. 2019. The centrosome protein AKNA regulates neurogenesis via microtubule organization. Nature. 567:113–117. doi:10.1038/s41586-019-0962-4.

Chuang, M., A. Goncharov, S. Wang, K. Oegema, Y. Jin, and A.D. Chisholm. 2014. The microtubule minus-end-binding protein patronin/PTRN-1 is required for axon regeneration in C. elegans. CellReports. 9:874–883. doi:10.1016/j.celrep.2014.09.054.

Cunha-Ferreira, I., A. Chazeau, R.R. Buijs, R. Stucchi, L. Will, X. Pan, Y. Adolfs, C. van der Meer, J.C. Wolthuis, O.I. Kahn, P. Schätzle, M. Altelaar, R.J. Pasterkamp, L.C. Kapitein, and C.C. Hoogenraad. 2018. The HAUS Complex Is a Key Regulator of Non-centrosomal Microtubule Organization during Neuronal Development. CellReports. 24:791–800. doi:10.1016/j.celrep.2018.06.093.

Efimov, A., A. Kharitonov, N. Efimova, J. Loncarek, P.M. Miller, N. Andreyeva, P. Gleeson, N. Galjart, A.R.R. Maia, I.X. McLeod, J.R. Yates, H. Maiato, A. Khodjakov, A.S. Akhmanova, and I. Kaverina. 2007. Asymmetric CLASP-dependent nucleation of noncentrosomal microtubules at the trans-Golgi network. Dev Cell. 12:917–930. doi:10.1016/j.devcel.2007.04.002.

Feng, C., P. Thyagarajan, M. Shorey, D.Y. Seebold, A.T. Weiner, R.M. Albertson, K.S. Rao, A. Sagasti, D.J. Goetschius, and M.M. Rolls. 2019. Patronin-mediated minus end growth is required for dendritic microtubule polarity. J Cell Biol. jcb.201810155–20. doi:10.1083/jcb.201810155.

Fernández, V., C. Llinares-Benadero, and V. Borrell. 2016. Cerebral cortex expansion and folding: what have we learned? EMBO J. doi:10.15252/embj.201593701.

Fietz, S.A., I. Kelava, J. Vogt, M. Wilsch-Bräuninger, D. Stenzel, J.L. Fish, D. Corbeil, A. Riehn, W. Distler, R. Nitsch, and W.B. Huttner. 2010. OSVZ progenitors of human and ferret neocortex are epithelial-like and expand by integrin signaling. Nat Neurosci. 13:690–699. doi:10.1038/nn.2553.

Guedes-Dias, P., J.J. Nirschl, N. Abreu, M.K. Tokito, C. Janke, M.M. Magiera, and E.L.F. Holzbaur. 2019. Kinesin-3 Responds to Local Microtubule Dynamics to Target Synaptic Cargo Delivery to the Presynapse. Curr Biol. 29:268–282.e8. doi:10.1016/j.cub.2018.11.065.

Hansen, D.V., J.H. Lui, P.R.L. Parker, and A.R. Kriegstein. 2010. Neurogenic radial glia in the outer subventricular zone of human neocortex. Nature. 464:554–561. doi:10.1038/nature08845.

Hu, D.J.-K., A.D. Baffet, T. Nayak, A.S. Akhmanova, V. Doye, and R.B. Vallee. 2013. Dynein recruitment to nuclear pores activates apical nuclear migration and mitotic entry in brain progenitor cells. Cell. 154:1300–1313. doi:10.1016/j.cell.2013.08.024.

Hu, W.F., O. Pomp, T. Ben-Omran, A. Kodani, K. Henke, G.H. Mochida, T.W. Yu, M.B. Woodworth, C. Bonnard, G.S. Raj, T.T. Tan, H. Hamamy, A. Masri, M. Shboul, M. Al Saffar, J.N. Partlow, M. Al-Dosari, A. Alazami, M. Alowain, F.S. Alkuraya, J.F. Reiter, M.P. Harris, B. Reversade, and C.A. Walsh. 2014. Katanin p80 regulates human cortical development by limiting centriole and cilia number. Neuron. 84:1240–1257. doi:10.1016/j.neuron.2014.12.017.

Jayaraman, D., B.-I. Bae, and C.A. Walsh. 2018. The Genetics of Primary Microcephaly. Annu Rev Genomics Hum Genet. 19:177–200. doi:10.1146/annurev-genom-083117-021441.

Jiang, K., S. Hua, R. Mohan, I. Grigoriev, K.W. Yau, Q. Liu, E.A. Katrukha, A.F.M. Altelaar, A.J.R. Heck, C.C. Hoogenraad, and A.S. Akhmanova. 2014. Microtubule Minus-End Stabilization by Polymerization-Driven CAMSAP Deposition. Dev Cell. 28:295–309. doi:10.1016/j.devcel.2014.01.001.

Kapitein, L.C., M.A. Schlager, M. Kuijpers, P.S. Wulf, M. van Spronsen, F.C. MacKintosh, and C.C. Hoogenraad. 2010. Mixed microtubules steer dynein-driven cargo transport into dendrites. Curr Biol. 20:290–299. doi:10.1016/j.cub.2009.12.052.

Kriegstein, A.R., and A. Alvarez-Buylla. 2009. The glial nature of embryonic and adult neural stem cells. Annu. Rev. Neurosci. 32:149–184. doi:10.1146/annurev.neuro.051508.135600.

Lui, J.H., D.V. Hansen, and A.R. Kriegstein. 2011. Development and evolution of the human neocortex. Cell. 146:18–36. doi:10.1016/j.cell.2011.06.030.

Marcette, J.D., J.J. Chen, and M.L. Nonet. 2014. The Caenorhabditis elegans microtubule minus-end binding homolog PTRN-1 stabilizes synapses and neurites. Elife. 3:e01637. doi:10.7554/eLife.01637.

Nakagawa, N., C. Plestant, K. Yabuno-Nakagawa, J. Li, J. Lee, C.-W. Huang, A. Lee, O. Krupa, A. Adhikari, S. Thompson, T. Rhynes, V. Arevalo, J.L. Stein, Z. Molnár, A. Badache, and E.S. Anton. 2019. Memo1-Mediated Tiling of Radial Glial Cells Facilitates Cerebral Cortical Development. Neuron. 103:836–852.e5. doi:10.1016/j.neuron.2019.05.049.

Noctor, S.C., A.C. Flint, T.A. Weissman, R.S. Dammerman, and A.R. Kriegstein. 2001. Neurons derived from radial glial cells establish radial units in neocortex. Nature. 409:714–720. doi:10.1038/35055553.

Noctor, S.C., V. Martínez-Cerdeño, L. Ivic, and A.R. Kriegstein. 2004. Cortical neurons arise in symmetric and asymmetric division zones and migrate through specific phases. Nat Neurosci. 7:136–144. doi:10.1038/nn1172.

Ori-McKenney, K.M., L.Y. Jan, and Y.-N. Jan. 2012. Golgi outposts shape dendrite morphology by functioning as sites of acentrosomal microtubule nucleation in neurons. Neuron. 76:921–930. doi:10.1016/j.neuron.2012.10.008.

Paridaen, J.T., and W.B. Huttner. 2014. Neurogenesis during development of the vertebrate central nervous system. EMBO Rep. 15:351–364. doi:10.1002/embr.201438447.

Pilaz, L.-J., A.L. Lennox, J.P. Rouanet, and D.L. Silver. 2016. Dynamic mRNA Transport and Local Translation in Radial Glial Progenitors of the Developing Brain. Curr Biol. 26:3383–3392. doi:10.1016/j.cub.2016.10.040.

Pinson, A., T. Namba, and W.B. Huttner. 2019. Malformations of Human Neocortex in Development - Their Progenitor Cell Basis and Experimental Model Systems. Front Cell Neurosci. 13:305. doi:10.3389/fncel.2019.00305.

Poirier, K., N. Lebrun, L. Broix, G. Tian, Y. Saillour, C. Boscheron, E. Parrini, S. Valence, B.S. Pierre, M. Oger, D. Lacombe, D. Geneviève, E. Fontana, F. Darra, C. Cances, M. Barth, D. Bonneau, B.D. Bernadina, S. N’Guyen, C. Gitiaux, P. Parent, V. des Portes, J.M. Pedespan, V. Legrez, L. Castelnau-Ptakine, P. Nitschke, T. Hieu, C. Masson, D. Zelenika, A. Andrieux, F. Francis, R. Guerrini, N.J. Cowan, N. Bahi-Buisson, and J. Chelly. 2013. Mutations in TUBG1, DYNC1H1, KIF5C and KIF2A cause malformations of cortical development and microcephaly. Nat Genet. 45:639–647. doi:10.1038/ng.2613.

Pollen, A.A., T.J. Nowakowski, J. Chen, H. Retallack, C. Sandoval-Espinosa, C.R. Nicholas, J. Shuga, S.J. Liu, M.C. Oldham, A. Diaz, D.A. Lim, A.A. Leyrat, J.A. West, and A.R. Kriegstein. 2015. Molecular identity of human outer radial glia during cortical development. Cell. 163:55–67. doi:10.1016/j.cell.2015.09.004.

Pongrakhananon, V., H. Saito, S. Hiver, T. Abe, G. Shioi, W. Meng, and M. Takeichi. 2018. CAMSAP3 maintains neuronal polarity through regulation of microtubule stability. Proc Natl Acad Sci USA. 115:9750–9755. doi:10.1073/pnas.1803875115.

Qu, X., A. Kumar, H. Blockus, C. Waites, and F. Bartolini. 2019. Activity-Dependent Nucleation of Dynamic Microtubules at Presynaptic Boutons Controls Neurotransmission. Curr Biol. 29:4231–4240.e5. doi:10.1016/j.cub.2019.10.049.

Reillo, I., C. De Juan Romero, M.Á. García-Cabezas, and V. Borrell. 2011. A role for intermediate radial glia in the tangential expansion of the mammalian cerebral cortex. Cerebral Cortex. 21:1674–1694. doi:10.1093/cercor/bhq238.

Reiner, O., R. Carrozzo, Y. Shen, M. Wehnert, F. Faustinella, W.B. Dobyns, C.T. Caskey, and D.H. Ledbetter. 1993. Isolation of a Miller-Dieker lissencephaly gene containing G protein beta-subunit-like repeats. Nature. 364:717–721. doi:10.1038/364717a0.

Sánchez-Huertas, C., F. Freixo, R. Viais, C. Lacasa, E. Soriano, and J. Lüders. 2016. Non-centrosomal nucleation mediated by augmin organizes microtubules in post-mitotic neurons and controls axonal microtubule polarity. Nature Communications. 7:12187–14. doi:10.1038/ncomms12187.

Shitamukai, A., D. Konno, and F. Matsuzaki. 2011. Oblique radial glial divisions in the developing mouse neocortex induce self-renewing progenitors outside the germinal zone that resemble primate outer subventricular zone progenitors. Journal of Neuroscience. 31:3683–3695. doi:10.1523/JNEUROSCI.4773-10.2011.

Tan, X., and S.-H. Shi. 2013. Neocortical neurogenesis and neuronal migration. Wiley Interdiscip Rev Dev Biol. 2:443–459. doi:10.1002/wdev.88.

Tas, R.P., A. Chazeau, B.M.C. Cloin, M.L.A. Lambers, C.C. Hoogenraad, and L.C. Kapitein. 2017. Differentiation between Oppositely Oriented Microtubules Controls Polarized Neuronal Transport. Neuron. 96:1264–1271.e5. doi:10.1016/j.neuron.2017.11.018.

Taverna, E., F. Mora-Bermúdez, P.J. Strzyz, M. Florio, J. Icha, C. Haffner, C. Norden, M. Wilsch-Bräuninger, and W.B. Huttner. 2016. Non-canonical features of the Golgi apparatus in bipolar epithelial neural stem cells. Sci Rep. 6:21206. doi:10.1038/srep21206.

Taverna, E., M. Götz, and W.B. Huttner. 2014. The cell biology of neurogenesis: toward an understanding of the development and evolution of the neocortex. Annu Rev Cell Dev Biol. 30:465–502. doi:10.1146/annurev-cellbio-101011-155801.

Tsunekawa, Y., J.M. Britto, M. Takahashi, F. Polleux, S.-S. Tan, and N. Osumi. 2012. Cyclin D2 in the basal process of neural progenitors is linked to non-equivalent cell fates. EMBO J. 31:1879–1892. doi:10.1038/emboj.2012.43.

Wang, Y., M. Rui, Q. Tang, S. Bu, and F. Yu. 2019. Patronin governs minus-end-out orientation of dendritic microtubules to promote dendrite pruning in Drosophila. Elife. 8:711. doi:10.7554/eLife.39964.

Wu, J., C. de Heus, Q. Liu, B.P. Bouchet, I. Noordstra, K. Jiang, S. Hua, M. Martin, C. Yang, I. Grigoriev, E.A. Katrukha, A.F.M. Altelaar, C.C. Hoogenraad, R.Z. Qi, J. Klumperman, and A.S. Akhmanova. 2016. Molecular Pathway of Microtubule Organization at the Golgi Apparatus. Dev Cell. 39:44–60. doi:10.1016/j.devcel.2016.08.009.

Yau, K.W., P. Schätzle, E. Tortosa, S. Pagès, A. Holtmaat, L.C. Kapitein, and C.C. Hoogenraad. 2016. Dendrites In Vitro and In Vivo Contain Microtubules of Opposite Polarity and Axon Formation Correlates with Uniform Plus-End-Out Microtubule Orientation. Journal of Neuroscience. 36:1071–1085. doi:10.1523/JNEUROSCI.2430-15.2016.

Yau, K.W., S.F.B. van Beuningen, I. Cunha-Ferreira, B.M.C. Cloin, E.Y. van Battum, L. Will, P. Schätzle, R.P. Tas, J. van Krugten, E.A. Katrukha, K. Jiang, P.S. Wulf, M. Mikhaylova, M. Harterink, R.J. Pasterkamp, A.S. Akhmanova, L.C. Kapitein, and C.C. Hoogenraad. 2014. Microtubule minus-end binding protein CAMSAP2 controls axon specification and dendrite development. Neuron. 82:1058–1073. doi:10.1016/j.neuron.2014.04.019.

Yokota, Y., T.-Y. Eom, A. Stanco, W.-Y. Kim, S. Rao, W.D. Snider, and E.S. Anton. 2010. Cdc42 and Gsk3 modulate the dynamics of radial glial growth, inter-radial glial interactions and polarity in the developing cerebral cortex. Development. 137:4101–4110. doi:10.1242/dev.048637.

Yokota, Y., W.-Y. Kim, Y. Chen, X. Wang, A. Stanco, Y. Komuro, W. Snider, and E.S. Anton. 2009. The adenomatous polyposis coli protein is an essential regulator of radial glial polarity and construction of the cerebral cortex. Neuron. 61:42–56. doi:10.1016/j.neuron.2008.10.053.

